# On growth and form of the distal air exchange surfaces within the lung

**DOI:** 10.1101/138974

**Authors:** David Warburton, Clarence Wigfall, Harvey Pollack, David Koos, Denise AlAlam, Wei Shi, Scott Fraser, Rex Moats

## Abstract

**Summary statement:** Employing novel imaging and computational approaches we here deduce some new concepts as to how the distal shaping of airway lineage stem and progenitor cells may contribute to the growth and form of the air exchange surface (AES) of the lung, distal to the bronchoalveolar duct junction (BADJ). We then propose that the AES extrudes from the sides and tips of distinct families of small ducts, apparently by buckling of the thinning luminal epithelium into the surrounding mesenchyme, stabilized firstly by rings of elastin and later by rings of collagen and elastin fibers, that surround the mouth of each distal unit of the AES. We propose this mechanism as a parsimonious solution to achieving the optimum form, packing density and functional efficiency of the AES, while allowing sufficient space between for the plumbing of conducting airways, pulsatile capillary blood vessels, lymphatics, nerves and other key matrix and cellular components within the interstitial mesenchyme.

**Abstract:** Herein we show, using several novel imaging and computational approaches, how the air exchange units (AEUs) of the lung develop from the tips and sides of distinct families of tortuous ducts, that themselves ramify as distinct families distal to the bronchoalveolar duct junctions (BADJs), prenatally in humans but postnatally in mice. The mature AEUs thus consist of indented spheroids tightly packed between quite regularly spaced distal ducts. Since the diameter of the BADJs and the distal ducts increases rather than decreasing during the formation of AEUs, we further deduce that the AEUs must form by circular epithelial precursor buckling at their mouths with reulsting extrusion of their lumen into the surrounding mesenchyme, stabilized firstly by interlocking rings of elastin and later by rings of elastin and collagen fibers surrounding the mouth of each of the prospective AEUs. Furthermore, we show that the surface of each of the AEUs is highly rugose, being indented by the capillary network that lies close beneath the AES membranes. We propose that, the kissing theorem proposed of Newton tha expresses the number of times billiard balls may touch within their frame is a parsimonious solution to achieving optimum packing of the distal AES unit spaces, while allowing sufficient space between them to allow for conducting airways, closely applied pulsatile capillary blood vessels, lymphatics, nerves and other key components of the interstitial mesenchyme.

## Introduction

The evolution of the lung gas exchange surface separated terrestrial animals from fish. Previously, on the basis of viewing histological sections as well as scanning electron micrographs, development of the distal air exchange surfaces (AES) of the lung was thought to arise by erection of septae that partition more primitive, pre-existing tubular canaliculae^1–7^. Herein, using several novel imaging approaches and computational postimaging space filling algorithms, together with new surface and volume rendering techniques, developed under the LungMAP consortium^8,9^, we can now view the structure and development of the AES from both a luminal as well as a volumetric space filling perspective. Based on these novel views, we now propose a modification of the theory of AES formation: that the AES appears to achieve the final form of small tightly packed, corrugated surface spheroids at least in large part by extrusion of the thinning and differentiating airway progenitor cell lined lumen outwards into the surrounding mesenchyme, both from the sides as well as from the tips of the distal airway ducts, at least five of which ducts arise below each bronchoalveolar duct junction and ramify outwards towards the pleura.

## Results and Discussion

We have previously reported, in LungMAP website^8^ as well as on the NIH Director’s Blog ^9^, but have not hitherto analyzed them in any great detail, novel interior views of the distal airways in fresh, hydrated lung tissues. These “fly down the airway” views were generated from microCT scan data of mature mouse lung using Lightwave imaging software to produce hyper-real views of the interior of branched airways. Analysis of these images show among many novel and interesting things that the terminal AES is comprised of spheroids that have a markedly corrugated internal surface (Fig 1 and accompanying movies). Meanwhile, a smooth tubular ring, fancifully resembling a rubber tyre inner tube, surrounds the neck of each prospective unit of the AES.

**Figure 1.**
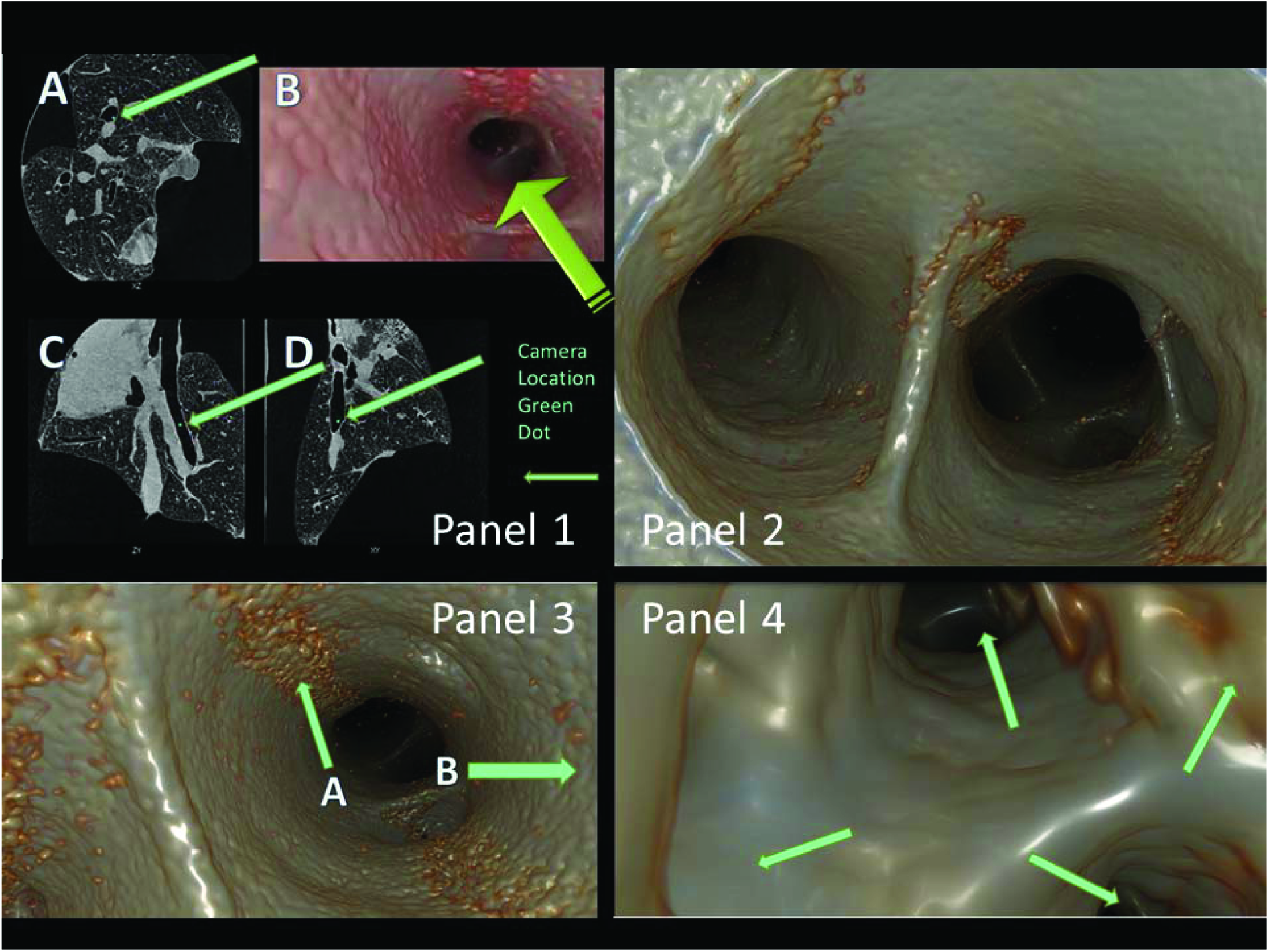
A “Fly down the airway” microCT Lightwave rendering of 28 day mouse lung, illustrating novel surface features of the airway and air exchange surface units of the lung. Panel 1. Shows microCT views of term mouse lung with position in the airway marked as a green dot of the camera view shown in B, which represents a bronchoalveolar duct junction (BADJ). Panel 2 shows a camera view distally from the carina down both major bronchi, which subdivide in the distance into lobar and sublobar bronchi. Novel features are the cobblestone appearance of Club cells in the epithelium and the curved shape of the carina and subsequent branch dividers. Panel 3 shows lumens of additional branches distal to the carina. The bumpy surface of the airway is seen to be caused by luminal cell surfaces fitting tightly like cobblestones into the epithelial surface. Also tracts of cilia coated with mucus are seen running down the sides of the airways A, B arrows. Panel 4. Shows the airway lumen distal to the bronchoalveolar duct junction (BADJ). The epithelia surface is now seen as smooth and flattened. Cilial tracts are no longer evident. The arrows indicate lateral and distal branches of the distal airway ducts (DADS). The green arrows indicate the smooth rounded mouths of sideways and distal branches of the DADS that proceed distally to terminate at air exchange surface units.

Space filling renderings of the AES have further revealed that the average mouth diameter of the terminal units of AES in adult mouse is 40 microns, and that they arise from the sides and tips of each of the 60-100 micron diameter distal airway ducts (DADS), 60 or more of which ramify from each 150 micron diameter bronchoalveolar duct junction (BADJ) (Figure 2 and accompanying movie). Moreover as the DADs extend outwards from the BADJs towards the pleura, their diameter remains relatively stable, while it increases rather than decreasing over the course of temporal expansion of the lung. Therefore it would seem to be a mathematical and physical impossibility that the AES can arise solely by erection or contraction of “septae” to subdivide pre-existing DAD spaces, neither in the mouse nor possibly in the human, no matter what viewing 2 dimensional sections may have appeared to suggest. It is also important to reciterate that the lung is continually expanding during the formation of the AES. Instead, therefore we are led to infer the new hypothesis that AES terminal lumens are extruded away from the DAD lumen, rather than being septated into it as has been previously widely accepted to be the case.

**Figure 2.**
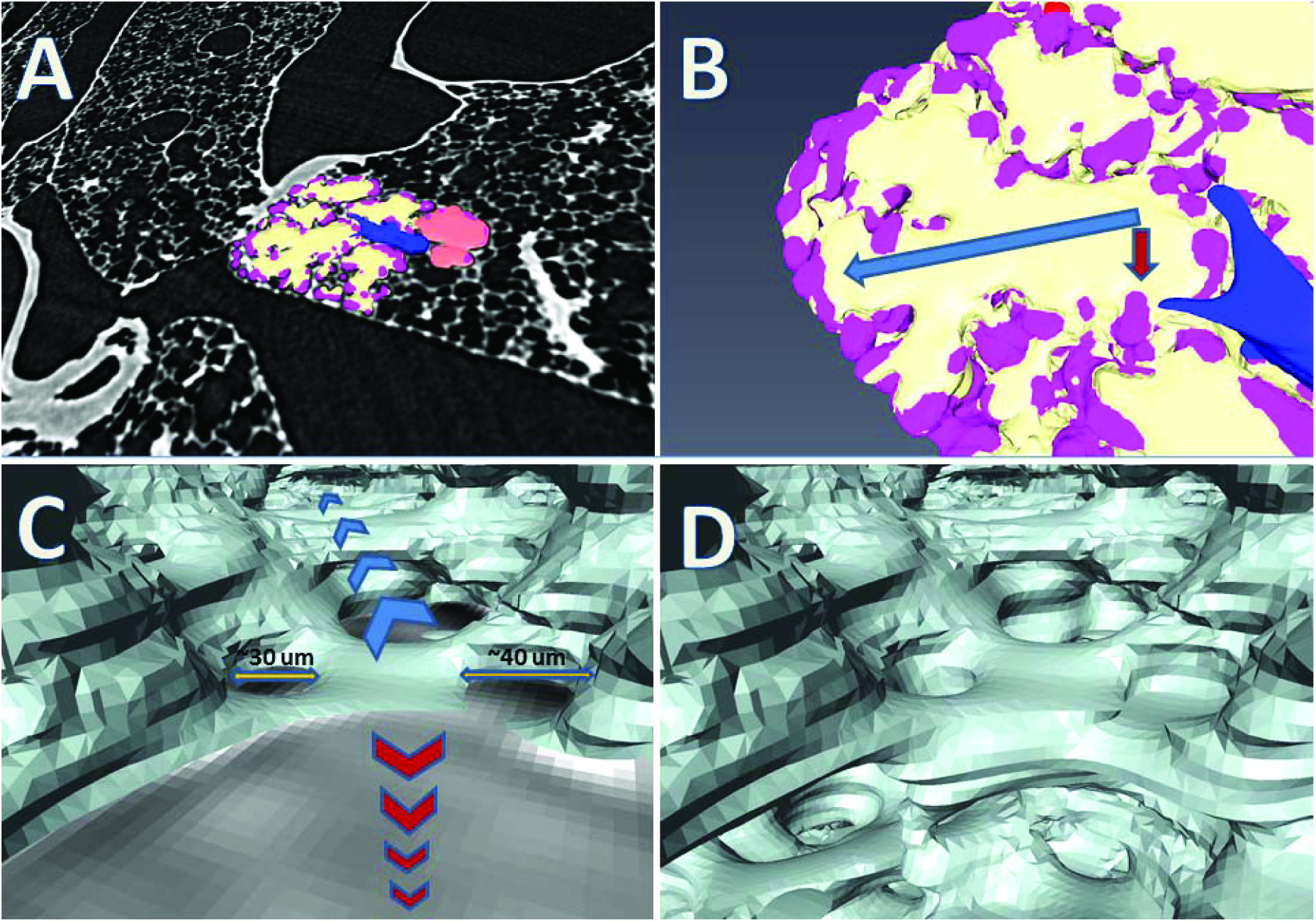
Space filling renderings illustrating novel features of the relationship, distal to the bronchoalveolar duct junctions, between distal airway ducts, DADs, the air exchange surface units and the aterial and capillary vasculature. A. Space filling rendering derived from microCT view of adult mouse lung. Yellow represents the air within a family of air exchange units surface units, orange represents air within a DAD dital to the BADJ. Blue represents deoxygenated blood retuning to the gas exchange surface down a distal pulmonary arteriole, which connects to an interlocking network of pulmonary capillaries shown in purple. B. Shows a magnified view of the space rendering in A. The color coding is the same. Arrows show the view trajectories of the yet more magnified perspective renderings in C and D. C. Rendered perspective view of the blue arrow in B. Each voxel in the figure represents 1 micron. Perspective effect is indicated by the changing sizes of the voxel sides. The view in the direction of the blue arrow disappear into the distance down a DAD towards the pleura. The floor is tinted red to provide an artificial scale plane. Note the round smooth 30-40 micron mouths of individual air exchange surface units that open into the DAD from both the sides and distally. D. The same view as in C, but with the artificial floor scale removed to allow simultaneous views at right angles along the paths of both the blue and red arrows in C showing how DADS can ramify at right angles distal to the BADJ to produce similar vistas of individual air exchange surface units arising perpendicular to the course of each DAD, and which do not encroach upon the DAD lumen, but arise outwards from the DAD lumen.

In addition we confirm, using volumetric modeling, that BADJs are established at least as early as E18.5 in mouse as well as long before birth in humans (Fig 3 and accompanying movies). Then, distal to the BADJs, at least 5 DADS extend from each BADJ and in many cases ramify at least 5 more times as far as the pleura, or in many other cases to the apparently limiting surface of interstitial fascial planes that subdivide the lung and bound individual non-overlapping families of AES units. We further find that circular, smooth rings comprising increased density of extracellular matrix proteins such as elastin and later collagen are formed along the sides as well as at the tips of individual DADS around the time of birth in mice, well prior to the appearance of any final form of the AES.

**Figure 3.**
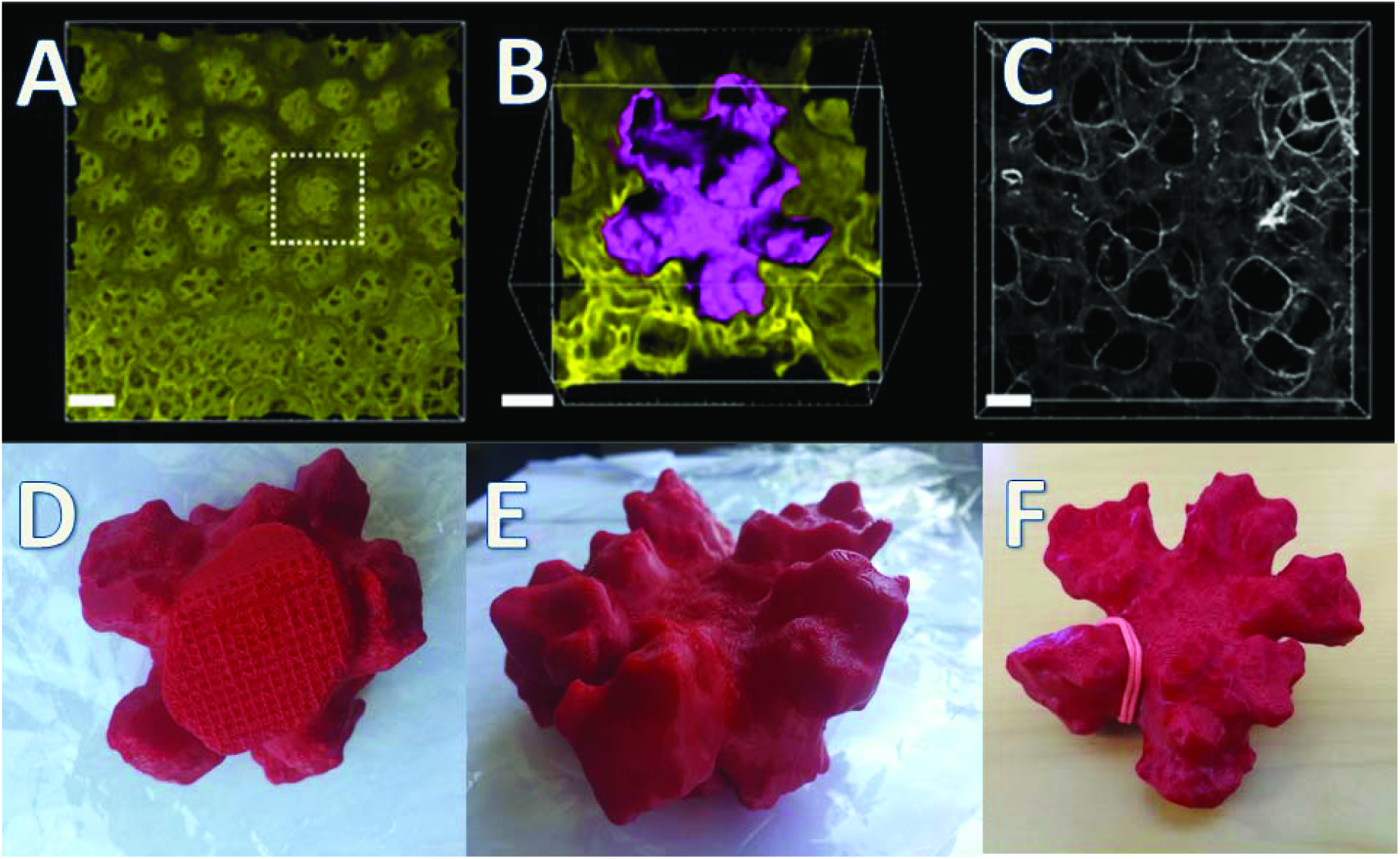
Distal families of air exchange surface (AES) units that arise from the tips of distal airway ducts in 28 day postnatal (adult) mouse lung. **A. Families each comprising several individual Units of the AES of the lung (box) lie at regular intervals just inside the pleura. The lung is taken from a membrane tomato mouse, sectioned, imaged by VIBRA-SSIM and the image reconstructed with software. At the entrance to each family of AES units can be seen a ring of tissue that surrounds the mouth of each family. Scale bar 50 microns.** **B. Surface rendering in purple and enlarged of the typical family of AES units shown in A. The surfaces of 5 major individual AES units are clearly interconnected as well having a strikingly rugose appearance. The surrounding mesenchyme cell membranes are rendered in gold in a sectional view. That there are many small vessels within the mesenchyme seen in cross section that indent the epithelial surface can be appreciated. The crisscross pattern of indentation appears to be rather similar to a Scots plaid with a gap between the stripes. Scale bar 15 microns.** **C. Autofluorescence image showing interlocking rings of collagen surrounding the mouth of each AES unit as well as surrounding the mouth of each family of units.** **D, E, F. Perspective views of three-dimensional printed volumetric rendering of the air exchange surfaces** shown above in Panel B, photographed from 3 different angles. D is a proximal to distal view similar to that in A. Note that the distal duct is 60 microns in diameter and from it subtend 5 major AES units. B. Note the striking rugosity of the air diffusion surface of each terminal unit in the family in this oblique view. F shows a distal to proximal plan view of the 5 major units when the rugosity of the surface can again be appreciated. The pale red elastic band was added to exemplify one of the collagen rings seen in C.

Another important observable feature, this time as expected, is that proximal to the BADJs, airway cells that line the lumen are mostly cuboidal in shape, with muffin like tops, whereas distal to the BADJs airway cells are much more flattened. This change in shape corresponds to differentiation of proximal versus distal epithelial progenitors into phenotypes of Club, mucus and ciliated cells proximal to the BADJ, whereas distal to the BADJ, Type 1 and 2 epithelial cells line the distal AES.

The idea that “septae” are erected by contractile elastin at their tips to subdivide pre-existing larger spaces or cannaliculae in the fetal lung in humans as well as the postnatal lung in mice has been repeated quite uncritically since well before 1970, including in many of our own papers and reviews. Now we come perforce with a seemingly heretical alternative hypothesis: that extrusion of lumens may be a more accurate way to describe the expansion of AES outwards from the sides and tips of DADS because DADS appear to retain or even enlarge their original diameter during the course of continuous expansion of the overall lung, and therefore cannot be subdivided by the erection of “septae”. This inference is based on volumetric rendering of the distal airways based on fresh microCT, as well as on pressure controlled perfusion fixed lungs using VIBRA-SSIM confocal approaches, with the aid of suitable advanced post image computational processing, including the use of Lightwave software borrowed from the movie industry (Figures 1–3).

The conclusions based on visualization techniques in the earlier literature such as histological sections and scanning EM were based on relatively harshly fixed, dehydrated specimens as well as inferences from the appearance of scanning EM or even Cyclotron images. These techniques may have introduced artefacts due to major shrinkage of tissues (>10%), which in turn results from the dehydration of the interstitium, as well as the deceiving perspective of the lateral, sectional point of view. Thus these previously established techniques may have given the erroneous impression that not only are capillary blood vessels collapsed, but also that the inferred septae are narrow with bulbous tips, that appear to be erected from their base outwards or upwards towards the lumen, rather than the lumen being extruded into the mesenchyme.

On other hand on the basis of our novel 3-dimensional geometry of hydrated as well as aqueous fixed fresh tissues, we now propose for the consideration of the community the alternative proposition that AES unit lumens actually extrude into the surrounding mesenchyme. As suggested in 1917 by Sir D’Arcy Wentworth Thompson in “On Growth and Form” ^15^, we here agree that surface tension must play a key role in forming soap films as well as similar thin walled lipid structures such as cell membranes and cells themselves. Thus, opening and stabilization of the AES unit lumens then appears to depend on differentiation and thinning of the distal epithelium, as well as on surfactant action to neutralize the tendency of small spheroid bodies to collapse, according to the Young-Laplace equation ^16–18^. Furthermore we suggest that the Navier-Stokes equations that govern the propagation of soap foams as well as the gas extrusion of bubbles into plastics may also apply here since AES may have several (as many as 3) stacked lumens^19^. Thus according to Thompson’s biomechanical model, in the intrauterine human fetal lung hydraulic fluid pressure must buckle and inflate the weaker and thinner distal AES membranous linings, with the matrix rings that surround their entrances serving as the stabilizing fulcrum for membrane buckling. Then we propose that distal cell membrane buckling is repeated, albeit on a smaller scale of the10 micron gap between the capillaries of the AES wall to give the plaid-like indentations of their walls. We then posit that small areas of the AES wall may also buckle and become extruded between a set of inward capillary indentations to form the next in a serial stack of AES lumens.

In the mouse as in the postnatal human the negative pressure of air breathing must be sufficient to drive this extrusion during further maturation of the AES in a process analogous to viscous fingering due to the Young’s modulus differential between air filled lumen of the epithelium and the adjacent hydrated mesenchyme. Even if we grant that the walls of individual AES units also do indeed grow in height, we interpret that since the proximal lumens to which they attach do not decrease in size, the direction of growth of the walls separating AES units must be outwards away from the 40 micron neck, thus extruding not subdividing the lumens as has previously been the accepted dogma, since the lung itself is also continuously expanding many fold distal to the BADJs during its development. We propose that this expansion, in conjunction with continual thinning of the intervening interstitial matrix space may be sufficient to explain how new units of AES can be accommodated within the limits of the continuously expanding lung “universe”.

In addition we propose that the progressive differentiation of epithelial progenitors distal to the BADJ into progressively thinner epithelial phenotypes must be important to allow buckling and extrusion of the distal AES membranes into the surrounding mesenchyme. The strain of buckling, extrusion and stretching of the epithelium also likely influences the adoption and organization of epithelial phenotypes. This latter idea is supported by single cell transcriptional analysis and genomic pathway analysis carried out in the LungMAP consortium and published on the limked websites LungGENS and Breath.

The astute reader will note that we have specifically eschewed using the relatively ancient ontologic term “alveoli” to describe the extruded peripheral AES of the lung. This intentional choice is suggested for possible adoption as common usage because “alveoli” has become conceptually encrusted over long usage, not only with incorrect epistomolgy: late 17^th^ Century derivation from Latin as a diminutive form of alveus, meaning a hollow trough-shaped vessel, trough, bath tub, hull, hold, ship, boat, channel or bed (of a river or trench) often containing something unpleasant such as excreta or rubbish, as well as by the concept of erection of “septae” into pre-existing air spaces. As Robert Hooke had it in 1664 in *Micrographia* ^20^, “By the addition of such artificial instruments and methods…and a wilfull and superstitious deserity of the Precepts and Rules of Nature,…every man is very subject to slip into all sorts of errors”. Therefore herein we respectfully suggest to the community the compromise usage air exchange surface (AES) units, which fortuitously overlaps with the existing ontologic term abbreviations AEC1 and AEC2, describing the two classes of epithelial cells lining the mature AES. We offer these new concepts about form and function of the air exchange surface within the lung for future discussion of its merits among the community, with rigorous experimental evaluation.

## Methods

Mouse lung preparation: All animal procedures and practices were approved by our institutional Animal Use and Care Committee. Before beginning the perfusion system is carefully degassed. The mouse was anesthetized to the surgical plane using intraperitoneal ketamine/xylazine. Next a tracheostomy was performed, the trachea intubated using a properly sized plastic catheter secured with an annular nylon suture. The catheter was then attached to a pressure controlled ventilator with settings appropriate to the animal weight and age, and ventilation with air started. Next a midline laparotomy was performed, the intestines retracted to expose the aorta and inferior vena cava (IVC), the IVC locally dissected free of the retroperitoneal tissue then cannulated with a plastic catheter secured with an annular nylon suture. Using a custom designed pressure controlled perfusion system an isotonic flush solution containing heparin was introduced through the IVC catheter then the aorta incised to allow blood to drain. Throughout the procedure perfusion pressure was actively maintained at 30 cm H_2_O. After the solution ran clear from the aortic incision airway pressure control was switched to a custom designed active airway pressure control system. The lung was “sighed” three times at a physiological rate cycling from 30 cm H2O to 10 cm H2O then the airway pressure was increased and maintained at 20 cm. Isotonic formaldehyde based fixative was then introduced through the IVC catheter until 60 ml of fixative had been passed into the mouse. Next the fixative flow was stopped, and a thoracotomy and anterior neck dissection performed. The heart, lungs, intact trachea and tracheostomy tube were removed en bloc while maintaining airway pressure. At this time specimens were either scanned immediately, transported to the microscopy suite for further processing, or dried at 35C for 5 days while the airway pressure is maintained.

MicroCT scan of the lungs: The lung specimens were scanned using the intact tracheostomy tube as the specimen mount. All scans were performed using a Bruker SkyScan 1172 (Bruker Inc. Billerica MA). Scan parameters were performed at the highest stable anode voltage possible of 60 kVp, with a tube current approximately 150 uA. Specific tube current was dependent on the age of the tube. 360 0.5 degree images were performed, with the detector fill factor of 70%. Scans took between 12 to 24 hours depending on specimen size. Reconstruction was performed using a Feldkamp filtered back projection type algorithm on software supplied by Bruker. Next the images were oriented to normal anatomic position and the right lobe of the lung segmented from the other tissue and background using manual segmentation on Amira Software (FEI Co, Hillsboro OR). After this the image stacks were converted to DICOM format using an application program written in-house, then stored on a PACS system.

The DICOM data was then accessed using LightWave animation software (NewTek Inc, San Antonio, TX). Using this software, volume rendering was completed and surface geometry applied to the data to create a 3D volume of the data as a high-resolution visual rendering. Using animation software to access the entire data set allowed any position within the lung rendering to be visualized. The animation software emulates camera and lighting positioning. The emulated camera was positioned within the upper trachea of the lung and directed on a virtual flight down through the airway all the way to the distal air exchange surface unit surface area. This camera movement was then rendered frame by frame as 4D animation with time being the 4^th^ dimension. This rendering was then formatted as 4K ultra high definition video suitable for use across the internet and for use directly from local hard drives and/or the cloud.

